# Face processing in the infant brain after pandemic lockdown

**DOI:** 10.1101/2022.01.26.477758

**Authors:** Tristan S. Yates, Cameron T. Ellis, Nicholas B. Turk-Browne

**Affiliations:** Department of Psychology, Yale University, New Haven, Connecticut, USA; Haskins Laboratories, New Haven, Connecticut, USA; Wu Tsai Institute, Yale University, New Haven, Connecticut, USA

**Keywords:** face identity, face recognition, experience, neuroimaging, fMRI

## Abstract

The role of visual experience in the development of face processing has long been debated. We present a new angle on this question through a serendipitous study that cannot easily be repeated. Infants viewed short blocks of faces during fMRI in a repetition suppression task. The same identity was presented multiple times in half of the blocks (Repeat condition) and different identities were presented once each in the other half (Novel condition). In adults, the fusiform face area (FFA) tends to show greater neural activity for Novel vs. Repeat blocks in such designs, suggesting that it can distinguish same vs. different face identities. As part of an ongoing study, we collected data before the COVID-19 pandemic and after an initial State lockdown was lifted. The resulting sample of 12 infants (9–24 months) divided equally into pre- and post-lockdown groups with matching ages and data quantity/quality. The groups had strikingly different FFA responses: pre-lockdown infants showed repetition suppression (Novel>Repeat), whereas post-lockdown infants showed the opposite (Repeat>Novel), often referred to as repetition enhancement. These findings provide speculative evidence that altered visual experience during the lockdown, or other correlated environmental changes, may have affected face processing in the infant brain.

## Introduction

What is the role of visual experience on the development of face processing? Although some researchers have argued that infants possess innate specialization for faces (Morton and Johnson, 1991; Turati, 2004; Kanwisher, 2010; Johnson et al., 2015), early visual experience can shape at least one aspect of face processing: the ability to distinguish individual faces (Pascalis et al., 2020). In a seminal study, 6-month-olds, 9-month-olds, and adults were shown pictures of human and monkey faces. All groups looked more at a novel human face than a familiar human face during test, evidence that they could distinguish human identities (Pascalis et al., 2002). However, only the youngest infants looked longer at a novel monkey face than a familiar monkey face, hinting that perceptual abilities narrow over early development (Maurer and Werker, 2014; Pascalis et al., 2020; Werker and Tees, 1984).

Evidence that perceptual narrowing is related to experience, rather than maturation, comes from studies that manipulate exposure to non-native faces (Sangrigoli and De Schonen, 2004; Pascalis et al., 2005; Anzures et al., 2012). For instance, infants exposed to monkey faces between 6 and 9 months are not only able to recognize the monkey faces to which they were exposed, but could also distinguish novel and familiar monkey faces from a new set when tested at 9 months (Pascalis et al., 2005). Even past 9 months, visual experience can influence the ability to process non-native faces, with infants recovering the ability to distinguish Asian female and male faces after daily exposure to videos of Asian women (Anzures et al., 2012). This period of plasticity is not indefinite: other-race (white) face recognition in Asian immigrants is significantly related to when in development they moved to a majority white country (Zhou et al., 2019).

Cases of deprivation provide an additional angle on the critical role that early visual experience plays in face identity processing. For instance, individuals with bilateral congenital cataracts at birth are impaired at holistic face processing (Le Grand et al., 2001; Grand et al., 2004) and individual face memory (de Heering and Maurer, 2014) even years after corrective surgery (Maurer 2017; cf. McKone et al. 2012). In another study, infant monkeys deprived of faces early in life were later shown either human or monkey faces for a month, prior to daily exposure to both species; immediately and one year later, monkeys could only distinguish face identities for the species to which they were initially exposed (Sugita, 2008; Arcaro and Livingstone, 2021).

It is unethical to conduct these kinds of causal tests in healthy human infants. In early 2020, however, the opportunity for a natural experiment in visual plasticity arose in response to the coronavirus disease 2019 (COVID-19) global pandemic. Emergency lockdowns and stay-at-home orders, as well as face mask coverings in public spaces, were used to prevent viral spread. These precautions abruptly changed daily life, including the nature of face-to-face interactions. Although infant face experience is highly biased toward primary caregivers (Sugden and Moulson, 2019), which continued and if anything may have expanded during lockdowns, absent or altered exposure to relatives and strangers during the COVID-19 pandemic may have influenced the development of face processing (Green et al., 2021; Carnevali et al., 2021). In particular, exposure to a more homogenous set of faces at home could alter identity processing. Indeed, smaller hometowns are associated with reduced accuracy on more difficult face recognition tasks (Balas and Saville, 2017; Sunday et al., 2019).

The tragic events of the COVID-19 pandemic therefore resulted in a quasi-experiment: we had been investigating neural measures of face identity processing in infants who underwent fMRI before the lockdowns were put in place, and then resumed data collection after a few months of the pandemic. Although the original purpose of this study was to investigate perceptual narrowing, as reflected in the design, we hypothesized that there may be a difference in face processing in the brains of infants tested pre- and post-lockdown. We used fMRI because it has the spatial resolution and sensitivity to resolve brain regions on the ventral surface of visual cortex distant from the scalp, including face-selective regions such as the fusiform face area (FFA). Recent studies established that the FFA is present in infants (Deen et al., 2017; Kosakowski et al., 2021). However, it is unknown whether the infant FFA additionally distinguishes face identities, similar to adult FFA, and if so, how the COVID-19 pandemic affects this processing.

Following a classic repetition suppression (or adaptation) design from adult fMRI (Grill-Spector et al., 2006; Turk-Browne et al., 2008), we showed infants blocks of faces that were of the same or different identities. In adults, the FFA tends to show reduced neural activity when the identity of a face is repeated vs. changed (Grill-Spector and Malach, 2001; Barron et al., 2016; Henson, 2016). We hypothesized from the start of this study that (pre-lockdown) infants would show repetition suppression in FFA. To the extent that the lockdown affected the development of face identity processing, we hypothesized that repetition suppression might be attenuated or eliminated. Specifically, if the post-lockdown infant FFA does not distinguish face identities, there should be no neural difference between same vs. different identities.

## Methods

### Participants

Data were collected from 12 infant fMRI sessions (9.20 – 23.80 months; *M* = 15.25, *SD* = 3.85; 9 female) that met inclusion criteria for 2 blocks of each condition, and with pairs of Novel and Repeat face blocks occurring in the same functional run (Table 1). Half of the sessions were collected prior to the onset of pandemic restrictions (until March 2020), and the other half of sessions were collected after our State’s first lockdown was lifted (from August 2020 until February 2021). As part of the original study, an additional 13 usable sessions were collected prior to the pandemic from infants under 9 months, but infants this young were not available post-lockdown for age matching purposes and so were not included in this analysis. We excluded data from 14 infant sessions in our target age range because an insufficient number of blocks were attempted or were retained after exclusion for eye movements or head motion. In the final sample, 2 unique infants provided 2 sessions of usable data each. Following prior work (Deen et al., 2017; Ellis et al., 2020; Kosakowski et al., 2021), the data from these sessions were treated as independent because they occurred several months apart (3.9 and 4.4 months). These second sessions were matched across groups, with one of the repeat infants in each of the pre- and post-lockdown groups, respectively. These infants saw a different set of images and counter-balancing in each session. The study was approved by the local Institutional Review Board. Parents provided informed consent on behalf of their child.

**Table 1.** Comparison of pre- and post-lockdown infant groups across demographic factors and measures of data quality and quantity. We found no reliable differences, all *ps* >0.1, with the exception of a marginally larger percent of TRs retained after exclusion for head motion for Repeat human blocks in pre-lockdown compared to post-lockdown infants.

|  | Pre-lockdown infants | Post-lockdown infants | *p*-value |
| --- | --- | --- | --- |
| N (female) | 6 (6) | 6 (3) | n/a |
| Days quarantine | n/a | 237.00 (52.91) | n/a |
| Age in months | 14.42 (3.19) | 16.08 (4.25) | 0.442 |
| Number blocks total | 13.33 (2.36) | 13.00 (2.16) | 0.732 |
| Percent gaze reliability | 95.6 (1.9) | 93.3 (4.3) | 0.214 |
| Percent TRs after motion (Novel Human) | 98.5 (3.3) | 97.1 (4.2) | 0.484 |
| Percent TRs after motion (Repeat Human) | 99.7 (0.7) | 95.6 (7.8) | 0.076 |
| Percent looking during exposure phase (Novel Human) | 89.5 (3.5) | 86.9 (7.7) | 0.524 |
| Percent looking during exposure phase (Repeat Human) | 88.4 (2.8) | 87.9 (4.9) | 0.860 |

### fMRI Data Acquisition

We followed validated procedures and parameters (see Figure 1A) for collecting fMRI data from awake infants (Ellis et al., 2020, 2021a,b,c; Yates et al., 2022). Data were acquired using the bottom half of a 20-channel head coil on a Siemens Prisma (3T) MRI. We collected functional images using a whole-brain T2* gradient-echo EPI sequence (TR = 2s, TE = 30ms, flip angle = 71, matrix = 64×64, slices = 34, resolution = 3mm iso, interleaved slice acquisition). For each session, we also collected anatomical images with a T1 PETRA sequence (TR1 = 3.32ms, TR2 = 2250ms, TE = 0.07ms, flip angle = 6, matrix = 320×320, slices = 320, resolution = 0.94mm iso, radial slices = 30,000).

**Figure 1.**
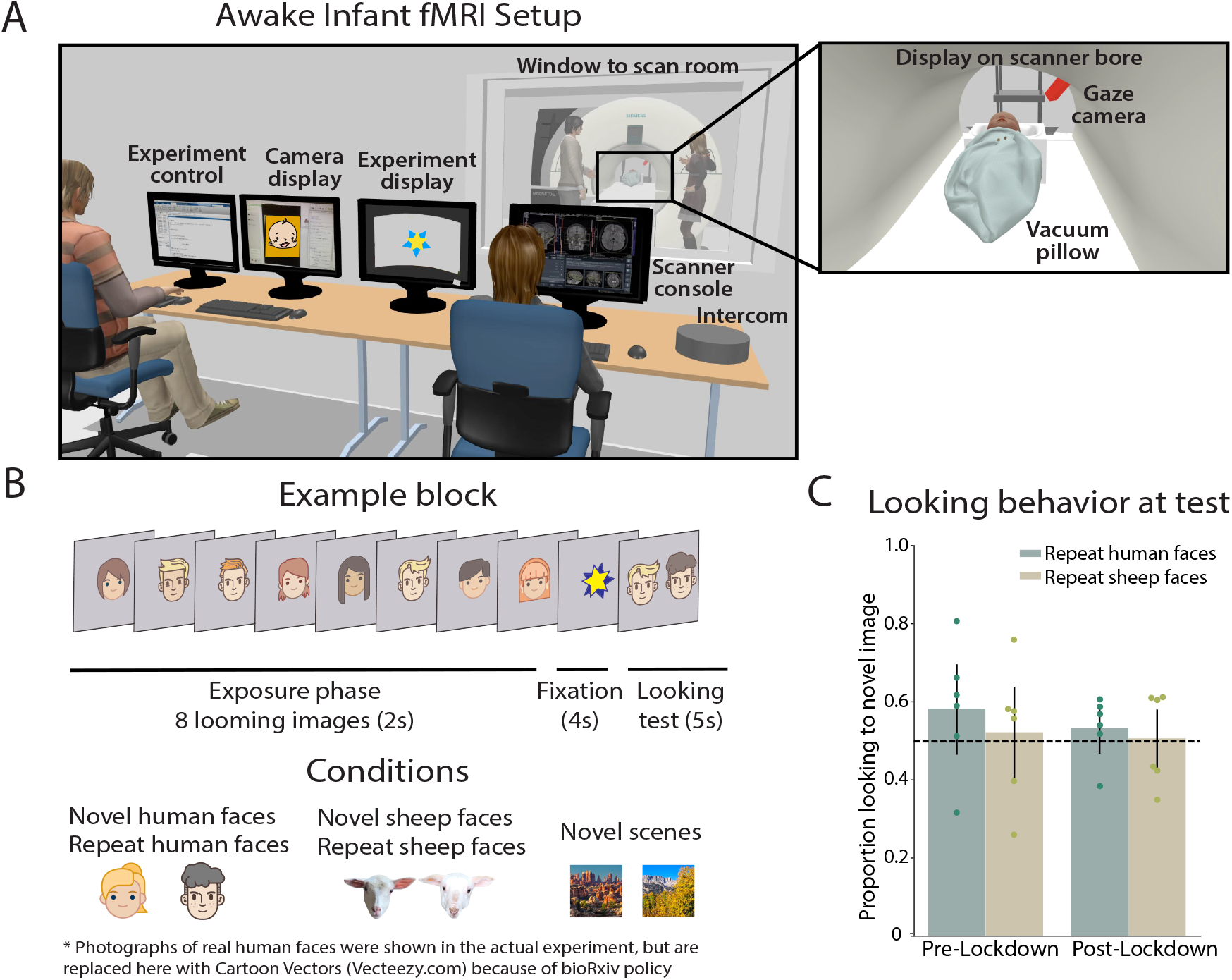
A. Setup for awake infant fMRI. In the control room, one experimenter monitors the infant and runs the tasks and another experimenter runs the scanner console and communicates with the experimenter in the scan room. In the scan room, an experimenter and parent stand on either side of the scanner bore. The infant is placed on a comfortable vacuum pillow at the center of the scanner bore with a panoramic view of the stimulus. A camera records the infant’s face. A full description of scanning methods is available (Ellis et al., 2020). B. During an experimental block, infants viewed a series of eight looming faces or scenes, followed by a short fixation period and a VPC looking test. Example images for different block conditions are shown (bottom). C. Infant behavior during the VPC test following Repeat face blocks for pre- and post-lockdown infants. There was no reliable evidence that infants in either group looked longer to the novel vs. repeated image, for either human or sheep faces. Dots represent individual participants.

### Procedure

Prior to their first scan, experimenters met with families for a mock scanning session. Families were invited for a scan session during a time when the parent thought the infant was most likely to be compliant. Before and at the scan visit, infants and accompanying parents were extensively screened for metal. Infants received three layers of hearing protection (silicon inner ear putty, over-ear adhesive covers, ear muffs) and were placed on top of a comfortable vacuum pillow on the scanner bed. We projected stimuli directly on the ceiling of the scanner bore above the infant’s face. We video recorded their face during scanning with a MRC high-resolution camera and coded their gaze offline. Procedures were identical for the pre-lockdown and post-lockdown groups, with the exception that for post-lockdown infants, all experimenters wore personal protective equipment for COVID-19 (respiratory mask, transparent face shield or goggles, and gloves), and parents wore masks throughout.

The task was presented to infants in MATLAB using Psychtoolbox (http://psychtoolbox.org/). Stimuli were human face images from the color FERET database (Phillips et al., 1998, 2000), outdoor scene images from an open dataset (http://olivalab.mit.edu/MM/sceneCategories.html; Konkle et al. 2010) and an internal database of web photos, and sheep face images (for the original perceptual narrowing study) photographed by the experimenters at a local sheep farm and supplemented through a web search. All stimuli were resized to 256 x 256 pixels. The background was cropped except for external features of the face (i.e., hair for humans, ears for sheep). Because fur color was consistent across sheep (all white/beige), we constrained the human face set to light-skinned or white human faces.

Each experimental block began with an exposure phase in which eight images were shown consecutively for two seconds each, looming from 1 to 20 visual degrees at the center of the screen (Figure 1B). An engaging attention-getter (rotating and expanding stars) was shown in the center of the screen for six seconds to encourage fixation. This was followed by a five-second visual-paired comparison (VPC) test phase, where one of the eight images from exposure was presented on one side and a novel image from the same category on the other side, separated by ten visual degrees. Each block thus lasted 27 seconds, followed by seven seconds of rest with a blank screen.

For the original purpose of the study, infants saw five block conditions: three were Novel blocks, in which each image was a new identity from the given category, and two were Repeat blocks, in which all eight images were the same identity. Scenes were only ever shown in Novel blocks, while human and sheep faces were shown in both Novel and Repeat blocks. Blocks were counter-balanced using a Latin-square design, with up to 25 blocks total available to participants. In a given session, we aimed to collect 2 blocks of each condition (10 total blocks = 5.7 minutes of data) but continued collecting more data if possible. Infants in the pre-lockdown group had 7.6 minutes (13.3 blocks) of usable data on average, and infants in the post-lockdown group had 7.4 minutes (13.0 blocks).

### Offline Gaze Coding

Infant gaze was coded offline by 2-3 coders who determined whether their eyes were looking center, right of center, left of center, off-screen (i.e., blinking or looking away), or undetected (i.e., out of the camera’s field ofview). Duringframes from the exposure and fixation phases, the coder was instructed that the infant was “probably looking at center.” This instruction was given to help calibrate coders to where the center of the screen was, but coders were allowed to indicate other responses if they believed the infant was not looking at the center. No instruction about likely looking was given for frames collected during the VPC test. Coders reported the same response code on an average of 94.4% (*SD* = 3.53%; range across participants = 83.8–97.6%) of frames. To combine across coders, we assigned each frame the modal response across coders from a moving window of five frames centered on that frame, and used the response from the previous frame in the case of ties. For a block to be included, infants needed to be looking at the screen (gaze coded as “center” during exposure and fixation phases, and either “left,” ‘right,” or “center” during VPC trials) for more than half of the frames.

We examined behavioral looking preference during the VPC test as the proportion of time looking to the novel image divided by the total time looking at either the familiar or novel image. The criteria for trials to be included in this analysis were that the infant looked at the familiar image during the earlier exposure phase (i.e., at center) and that they attended to the VPC test (i.e., at left, right, or center) for at least 500 milliseconds. We only analyzed behavior during VPC tests for Repeat blocks that were included in the fMRI analyses. We used non-parametric bootstrap resampling (Efron and Tibshirani, 1986) to test for significant preferences to the novel image. Proportion looking to the novel image was first averaged within a subject for a given condition. We then sampled average participant data from each group with replacement 1,000 times, calculating the average looking preference on each iteration. We calculated the *p*-value as the proportion of samples for which the mean was in the opposite direction from the true effect, doubled to make the test two-tailed.

### fMRI Preprocessing

We preprocessed data using a pipeline that has been used in prior awake infant fMRI studies and released publicly (Ellis et al., 2020). We sometimes collected tasks for other studies in the same functional runs. When this occurred (*N* = 9), the data were separated into pseudoruns for each task. In two sessions, experimental blocks were separated by a long break within session (205 and 2,913 seconds). Otherwise, participants viewed all blocks within the same functional run.

We removed three burn-in volumes from the beginning of each run/pseudorun. The centroid volume of each run/pseudorun (i.e., the volume that minimized the Euclidean distance to all other volumes) was used as the reference volume for motion correction and anatomical alignment. Slice-timing correction was used to realign slices in each volume. We excluded timepoints with greater than 3mm of translational motion; across participants, the majority of timepoints were included after motion exclusion (*M* = 97.3%, *SD* = 3.9%; range across participants = 86.6–100%). Excluded time points were interpolated to not bias the linear detrending, and then ignored in later analyses. We also excluded blocks of data if more than 50% percent of the timepoints were excluded due to motion or infants looking away from the screen. The mask of brain vs. non-brain voxels was formed by calculating the signal-to-fluctuating-noise ratio (SFNR) (Friedman and Glover, 2006) for voxels in the centroid volume. A Gaussian kernel (5mm FWHM) was used to spatially smooth the data. Data were also linearly detrended in time, and aberrant timepoints were attenuated using AFNI’s (https://afni.nimh.nih.gov) despiking algorithm.

We registered the centroid volume for each run/pseudorun to the infant’s anatomical image. Alignment was initially performed using FLIRT with a normalized mutual information cost function and six degrees of freedom (DOF). After manual inspection, this automatic registration was corrected if necessary using mrAlign from mrTools (Gardner lab). Functional data were then transformed into standard adult MNI space to make comparisons across infants. First, functional data were linearly aligned to an age-specific infant template using 12 DOF. This alignment was improved with non-linear warping using diffeomorphic symmetric normalization (ANTS; Avants et al. 2011). A predefined transformation (12 DOF) between the infant template and adult standard was then used. For all analyses, we only considered voxels included in the intersection of all infant brain masks.

### Regions of Interest

Regions of interest (ROIs) were defined using Neurosynth, a meta-analytic tool that combines results from published fMRI studies on different topics (Yarkoni et al., 2011). We used the search term “face” and obtained a whole-brain statistical map showing *z*-scores from a two-way ANOVA testing the presence of activated voxels associated with this term. This map indicates which regions are more consistently activated in the 896 studies about faces compared to all other studies in the database. The resulting map was thresholded using a false discovery rate of 0.01. The coordinates for peak activation in the anatomical vicinity of the right and left FFAwere used as the centers of two 10mm radius spheres around these peaks of activation. This bilateral FFA mask was used as the search space for our leave-one-participant-out functional ROI (fROI) analysis. We also created spheres around the peak activations for other regions known to be involved in face processing (Haxby et al., 2000): bilateral occipital face area (OFA), bilateral superior temporal sulcus (STS), bilateral amygdala (Amyg) and right inferior frontal gyrus (rIFG). Finally, we used the same procedure with search terms “primary visual cortex” and “heschl’s gyrus” to obtain whole brain statistical maps for defining control regions in primary visual cortex and primary auditory cortex, respectively.

### Statistical analyses

We fit general linear models (GLMs) to preprocessed BOLD activity using FEAT in FSL. Data were split and *z*-scored within functional runs according to three different GLM analyses: (1) a balanced number of Novel human face blocks and scene blocks, (2) a balanced number of Novel and Repeat human face blocks, and (3) a balanced number of Novel and Repeat sheep face blocks. In each GLM, regressors were created forthe 16-second exposure phases of each of the two modeled conditions, as well as for 6-second fixation periods and 5-second VPC tests combined across the two modeled conditions (to isolate exposure differences). Events were modeled using a boxcar for their duration, convolved with a double-gamma hemodynamic response function (Deen et al., 2017; Ellis et al., 2020). Motion parameters (3 translation and 3 rotation) from motion correction were included in the GLM as regressors of no interest. Time points excluded for high motion were scrubbed with an additional regressor for each time point.

The main contrast of interest for the first GLM was Novel human faces greater than scenes during the exposure phase, to reveal face-selective visual responses. Forthe second GLM, the contrast of Novel greater than Repeat human faces during the exposure phase provided an index of identity processing, with positive values reflecting repetition suppression. The third GLM tested for repetition suppression of sheep faces. The *z*-statistic volumes from these contrasts were extracted for each participant and aligned to standard space.

Our first objective was to determine whether an infant-defined FFA showed repetition suppression to Novel vs. Repeat human face blocks, and whether this differed across the pre- and post-lockdown groups. To accomplish this, we extract neural responses using an fROI approach (Figure 2A). From the first GLM and collapsing across pre- and post-lockdown groups (to increase power and avoid bias), we averaged the contrast of Novel human faces greater than scenes in standard space from all but one (held-out) infant. Using the meta-analysis FFA sphere as a mask, we located the top 5% of voxels that showed the greatest average contrast value (i.e., face selectivity; Figure 2B). Retaining these voxels, we then extracted the activation to Novel and Repeat human face blocks from the second GLM of the held-out infant, and averaged across voxels within condition. This procedure was iterated such that each infant was held out once. Critically, the fROI was defined independently of the data from the held-out infant to prevent circularity.

**Figure 2.**
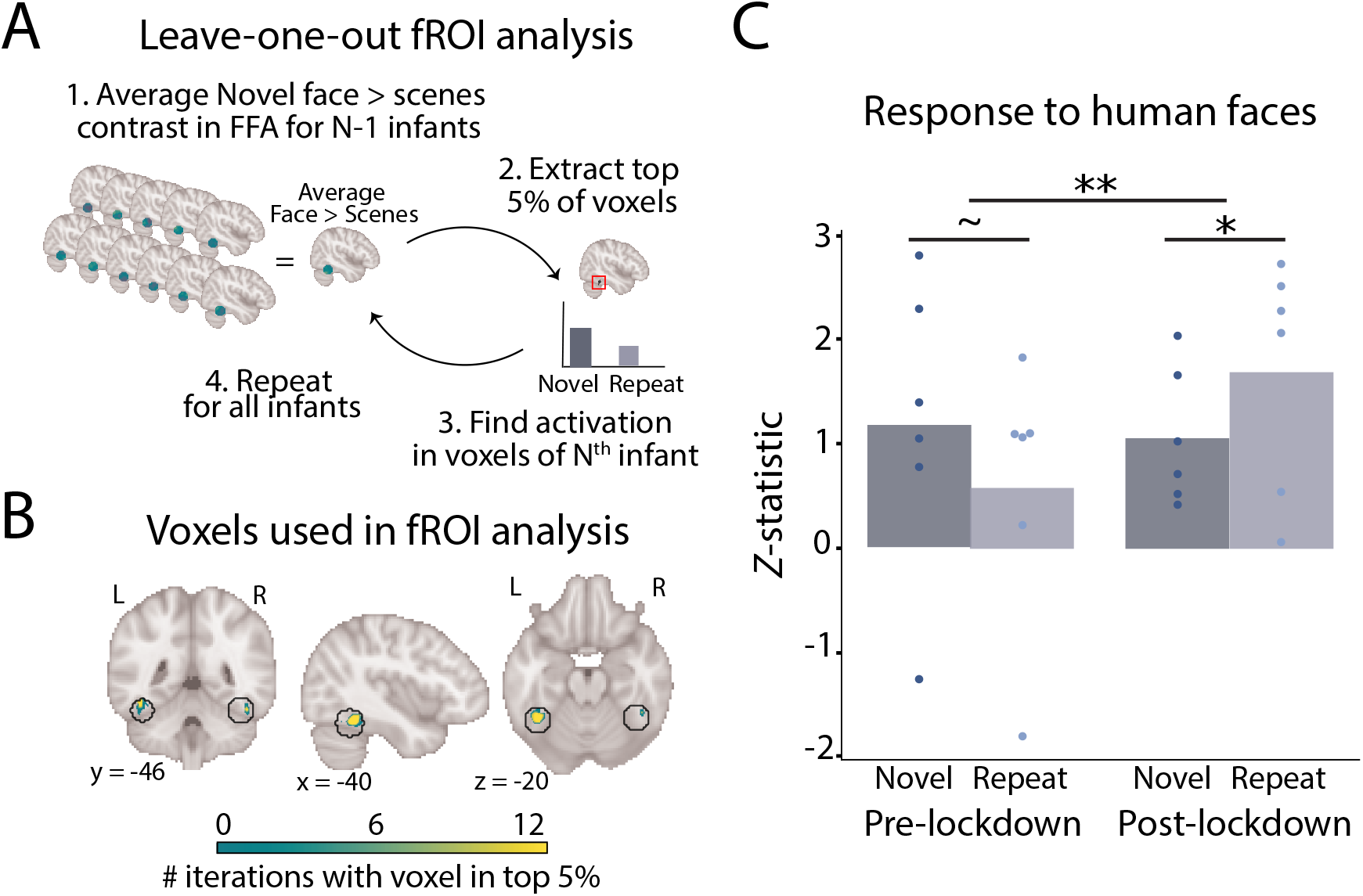
A. Leave-one-out fROI analysis. First, we averaged the *z*-statistics for the contrast of Novel human face blocks greater than scene blocks in *N* minus one infants. We selected the voxels with the top 5% of average values as being the most face-selective to define an fROI independently of the remaining infant. We then extracted the average *z*-statistics for Novel and Repeat human face blocks from these voxels in this held-out infant. We repeated the procedure 12 times so that each infant was held-out once. B. Voxels used in the fROI analysis. The circles outline the spherical FFA search space from the meta-analysis. Each voxel is colored by the number of iterations (of 12) in which it was among the top 5%. C. There was a robust group difference in the contrast of Novel vs. Repeat human face blocks: the neural response was marginally greater for Novel than Repeat human face blocks in pre-lockdown infants (repetition suppression), and significantly greater for Repeat than Novel in post-lockdown infants (repetition enhancement). Dots are individual participants. ** *p* < 0.01, * *p* < 0.05, ~*p* < 0.10.

We used non-parametric bootstrap resampling to test for the statistical significance of the extracted fROI data. For each infant group and condition, we resampled *z*-statistic values for the contrast of Novel vs. Repeat blocks (our measure of repetition suppression) with replacement 1,000 times and recalculated the average for each iteration. The *p*-value was then the proportion of resamples that were of the opposite sign as the original effect, doubled for a two-tailed test. We similarly quantified group differences in the contrast of Novel vs. Repeat by performing 1,000 resamples of *z*-statistic contrast values for the pre-lockdown and post-lockdown groups. On each iteration, we recalculated the mean value for each group and then subtracted the post-lockdown mean from the pre-lockdown mean. Again, the *p*-value was the proportion of resamples that were of the opposite sign as the original effect, doubled for a two-tailed test.

We next assessed whether our results generalized beyond this fROI approach by extracting the *z*-statistic values for Novel and Repeat human face blocks in all voxels from the meta-analysis FFA sphere (i.e., not just the top 5%) and averaged within condition. We also measured responses in the other meta-analysis face ROIs and control ROIs to test specificity to the FFA.

Finally, we assessed whether group differences were specific to human faces, or might apply more broadly to identity processing of other types of faces by repeating these analyses for the sheep data.

### Code and data availability

The code for the task is available at: https://github.com/ntblab/experiment_menu/tree/RepetitionNarrowing. The code for the analyses is available at: https://github.com/ntblab/infant_neuropipe/tree/RepetitionNarrowing. Raw and preprocessed functional and anatomical images will be released publicly upon acceptance

## Results

### Behavioral looking time to novel images

We first asked whether the pre- and post-lockdown infants differed in their amount of looking to novel vs. familiar faces. We focused on VPC tests that followed Repeat blocks because these blocks contained repetitions that might habituate infants to a face identity, allowing them to make a discrimination between the novel and familiar test items. There was no group difference in novelty preferences for human faces (*M* = 0.050, bootstrap *p* = 0.466; Figure 1C), with neither group showing a reliable preference relative to baseline (pre-lockdown: proportion looking to novel *M* = 0.584, CI = [0.465, 0.697], vs. 0.5 *p* = 0.162; post-lockdown: *M* = 0.534, CI = [0.468, 0.584], *p* = 0.278). This may in part reflect the relatively short habituation time and the use of multiple identities and blocks across the study. Likewise, there was no group difference in novelty preferences for sheep faces (*M* = 0.016, *p* = 0.848), and again neither group showed a reliable preference relative to baseline (pre-lockdown: *M* = 0.522, CI = [0.396, 0.639], *p* = 0.726; post-lockdown: *M* = 0.506, CI = [0.419, 0.581], *p* = 0.890). This is more expected given research showing a decline in discrimination of other-species faces after 9 months (Pascalis et al., 2002; Simpson et al., 2011).

### Neural responses to human face identity in infant FFA

We next asked whether the pre- and post-lockdown infants differed in their neural responses to human faces. There was a significant group difference in the FFA fROI for the contrast of Novel vs. Repeat human face blocks (z-score *M* = 1.499, bootstrap *p* = 0.006; Figure 2C). In the pre-lockdown group, there was marginally greater activation in the FFA fROI for Novel blocks (z-score *M* =1.185) than Repeat blocks (*M* = 0.588; difference *M* = 0.749, CI = [−0.096, 1.648], *p* = 0.088), consistent with the repetition suppression hypothesized for this group. In the post-lockdown group, however, there was *less* FFA fROI activation in Novel blocks (*M* = 1.067) than Repeat blocks (*M* = 1.705; difference *M* = −0.749, CI = [−1.617, 0.016], *p* = 0.033). This significant repetition *enhancement* effect implies that infants in the post-lockdown group were still able to neurally distinguish face identities, but responded differently to the repetitions.

The group difference in the contrast of Novel vs. Repeat human face blocks remained significant when considering the entire spherical FFA ROI from the meta-analysis of adult studies, rather than constraining it to face-selective voxels from (other) infants (*M* = 1.325, *p* = 0.004; Figure 3B). In pre-lockdown infants, the pattern was similar to the FFA fROI (Novel >Repeat) but not significant (*M* = 0.574, CI = [−0.136, 1.282], *p* = 0.124). In post-lockdown infants, the difference observed with the fROI approach (Repeat >Novel) remained significant (*M* = −0.751, CI = [−1.337, −0.207], *p* = 0.004). Thus, our findings were fairly robust to the number and selectivity of voxels contained in the FFA.

**Figure 3.**
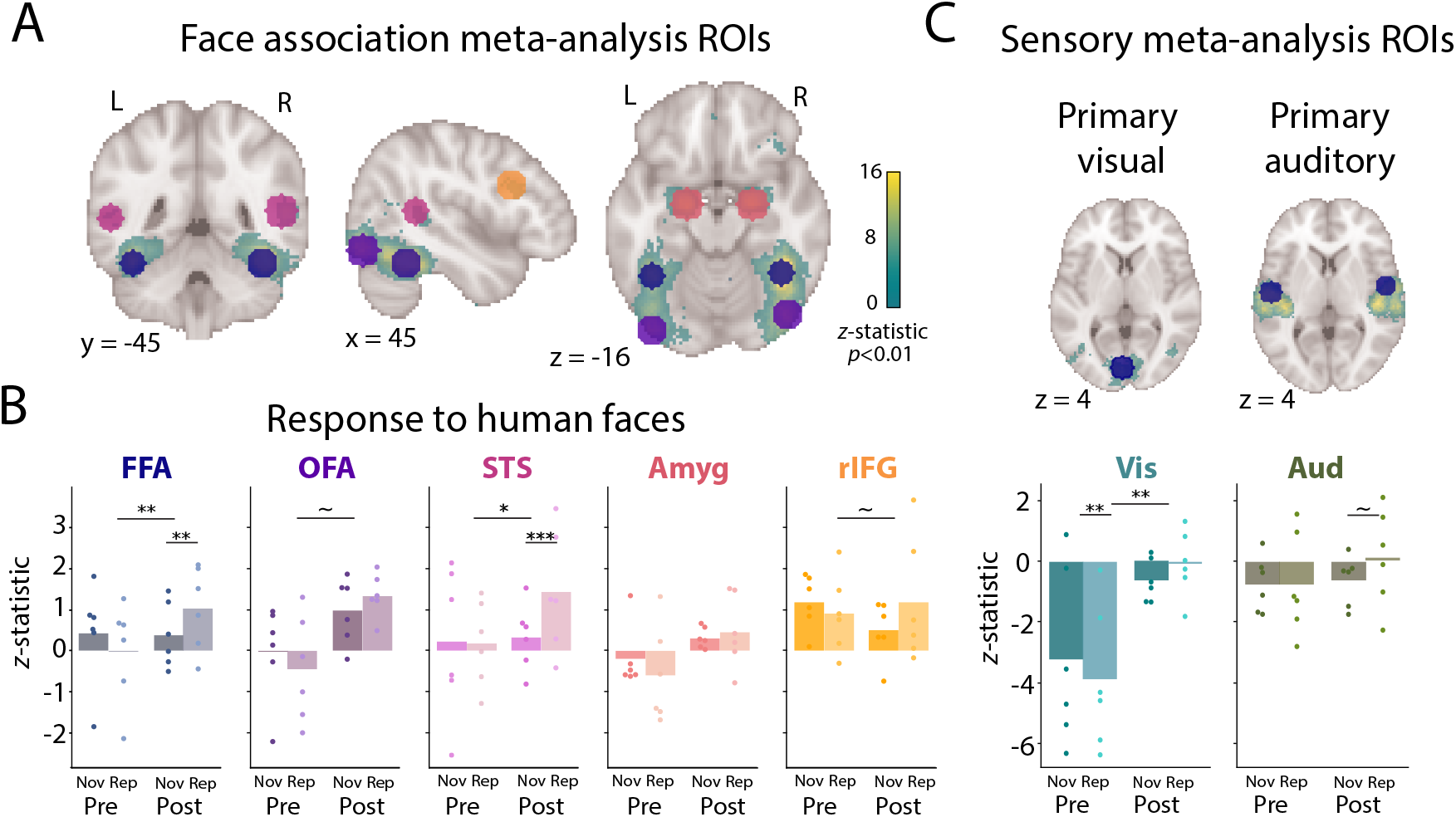
A. Spherical ROIs derived from an adult meta-analysis for the term “face” (Yarkoni et al., 2011). The voxel with the peak *z*-statistic value was assigned the center of a 10mm radius sphere. B. The spherical ROI containing all FFA voxels largely mirrored the results from the FFA fROI limited to the most face-selective voxels. A similar group difference in the Novel vs. Repeated contrast was found in the spherical ROI for STS and marginally for OFA and rIFG, but not for amygdala. C. The spherical ROI in primary visual cortex but not primary auditory cortex showed a similar group difference. The overall negative activity in primary visual cortex may reflect the inclusion of voxels responsive to the periphery of the visual field that did not contain stimuli. Dots are individual participants. *** *p* < 0.001, ** *p* < 0.01, * *p* < 0.05, ~*p* <0.10. ROIs: fusiform face area (FFA), occipital face area (OFA), superior temporal sulcus (STS), amygdala (Amyg) right inferior frontal gyrus (rIFG), primary visual cortex (Vis), and primary auditory cortex (Aud)

### Specificity of findings to FFA

We next investigated the specificity of the key group difference in facial identity processing to the FFA by considering spherical ROIs in the broader face processing network (OFA, STS, amygdala, rIFG; Figure 3A). The group difference between pre- and post-lockdown infants in the Novel vs. Repeat contrast was significant in the STS (*M* = 1.251, *p* = 0.018) and marginal in OFA (*M* = 0.946, *p* = 0.058) and rIFG (*M* = 1.089, *p* = 0.083), but not in amygdala (*M* = 0.670, *p* = 0.157; 3B). This effect extended to a spherical ROI in primary visual cortex (*M* = 1.372, *p* = 0.009) but not primary auditory cortex (*M* = 0.839, *p* = 0.218; Figure 3C). If correcting for multiple comparisons across all seven spherical ROIs, only the FFA survives (Bonferroni *p* = 0.007). These results indicate relative specificity of the group effect to the FFA.

### Specificity of findings to human faces

Finally, we examined the specificity of the results to human faces by repeating the analyses above for sheep face blocks (Figure 4). We did not use the FFA fROI because the voxels in the fROI were chosen based on responses to human face blocks, which might introduce a bias in favor of specificity to human faces. However, considering all voxels in the spherical ROI for FFA, there was a marginal group difference in the contrast of Novel vs. Repeat sheep face blocks (*M* = 1.067, *p* = 0.054), which did not survive correction for multiple comparisons. There was a greater neural response for Novel than Repeat sheep face blocks in pre-lockdown infants (*M* =1.120, CI = [0.622, 1.625], *p* <0.001) but no effect in either direction in post-lockdown infants (*M* = 0.052, CI = [−0.702, 1.013], *p* = 0.962). Thus, FFA results for sheep faces were similar to human faces, with a weaker group difference driven by the lack of sensitivity to sheep face identity in post-lockdown infants who had shown strong repetition enhancement for human face identity.

**Figure 4.**
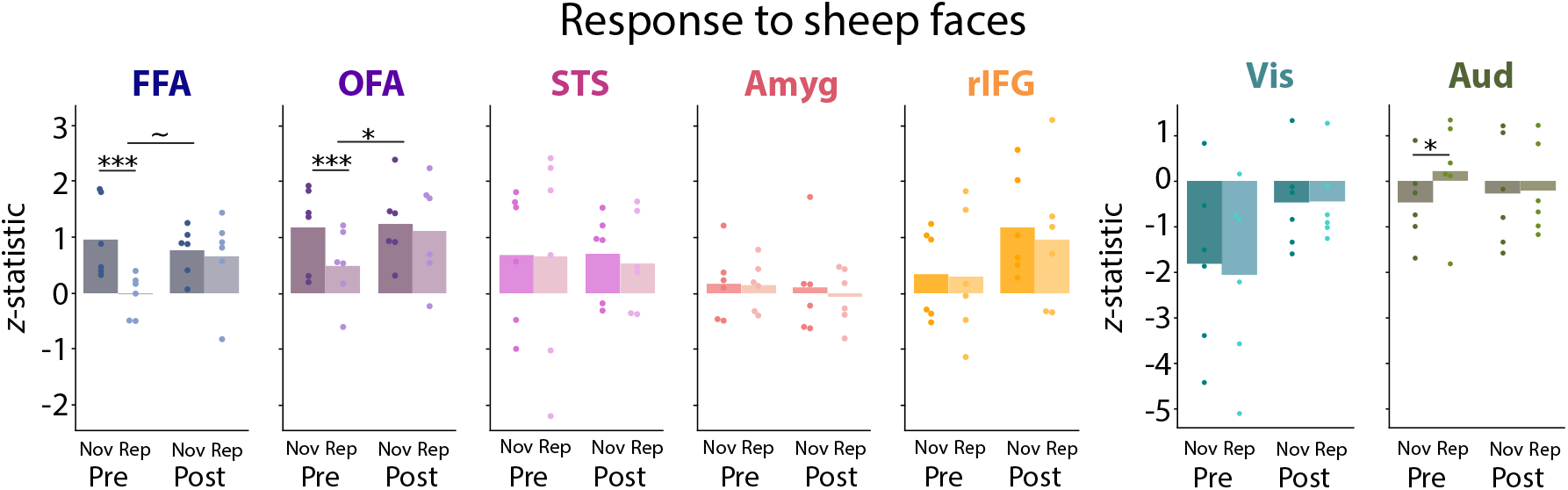
There was a group difference in the Novel vs. Repeat contrast for sheep faces in the spherical ROI for OFA and marginally for FFA, butwas not for STS, amygdala, rIFG, primary visual cortex, or primary auditory cortex. Dots represent individual participants. *** *p* < 0.001, * *p* < 0.05, ~*p* < 0.10.

When considering sheep face blocks in the broader set of spherical ROIs for face and sensory processing, we again focused on the key group difference in the contrast of Novel vs. Repeat. Only the OFA showed a significant group difference (*M* = 0.751, *p* = 0.049, which did not survive multiple comparisons correction. The other spherical ROIs did not show a reliable group difference: STS (*M* = −0.098, *p* = 0.837), amygdala (*M* = −0.154, *p* = 0.695), rIFG (*M* = −0.125, *p* = 0.836), primary visual cortex (*M* = 0.195, *p* = 0.699), and primary auditory cortex (*M* = −0.733, *p* = 0.138).

## Discussion

We investigated face identity processing in the brains of infants tested before and after the initial COVID-19 lockdown in our State. The neural responses of these groups to human faces differed: pre-lockdown infants showed some evidence of repetition suppression (Novel >Repeat) for human faces in the FFA, similar to what is seen in adults (Grill-Spector and Malach, 2001) and older children (Natu et al., 2016); post-lockdown infants showed the opposite, repetition *enhancement* (Repeat >Novel). This group difference was most robust in the FFA compared to other brain regions and for human faces compared to sheep faces.

This pattern of results is reminiscent of the debate over novelty vs. familiarity preferences in infant behavior (e.g., reduced vs. increased looking at repeated stimuli). Familiarity preferences are more likely in younger infants, for more complex stimuli, after shorter exposure, and for more difficult tasks (Rose et al., 1982; Hunter and Ames, 1988; Roder et al., 2000). Likewise, the adult brain shows increased processing of repeated stimuli when the stimuli are unfamiliar or briefly exposed (Henson et al., 2000; Turk-Browne et al., 2007; Segaert et al., 2013). By this account, human faces could be considered a more familiar category to pre-lockdown infants, with exposure to a greater number of unique faces buildinga more robust face space (Humphreys and Johnson, 2007), than to post-lockdown infants with more restricted face experiences. In particular, the pandemic presumably reduced exposure to faces that would have been experienced infrequently (e.g., relatives and strangers). This lends credence to the possibility that even limited exposure to different face exemplars can impact face processing (Spangler et al., 2013).

Notably, our behavioral measure in the scanner did not mirror the neural measure, as might have been expected (Snyder and Keil, 2008; Turk-Browne et al., 2008; Nordt et al., 2016). We interpret these null behavioral results with caution given the difficulty of collecting reliable behavior in the scanner even in adults and the possibility of different sensitivity and noise for neural and behavioral measures. Nevertheless, adult fMRI studies have shown that repetition suppression and enhancement can occur in the absence of (Segaert et al., 2013), and be dissociated from (Xu et al., 2007), behavioral measures of priming. Moreover, the similar pattern of looking behavior between groups during exposure and test phases is inconsistent with an attentional explanation of the group difference in neural responses, whereby the strength of neural responses might be an artifact of the amount of stimulus viewing. We believe that our findings illustrate the benefit of using multiple measures to study infant cognition (LoBue et al., 2020) and the potential of brain imaging to disentangle cognitive processes (Yates et al., 2021).

It is tempting to interpret these data in relation to how pandemic precautions such as social distancing and face masks altered early face experience, especially given the strict local guidelines that were imposed. However, this link is speculative in our study because we did not measure daily exposure to faces in the pre- or post-lockdown group; indeed, this study was designed and partially completed prior to the pandemic. Thus, we cannot conclude definitively that our results are related to altered visual experience with faces. The pandemic had many other impacts on daily life that could explain neural differences. Perhaps most relevant to face processing is an increase in maternal fear and anxiety (Cameron et al., 2020; Davenport et al., 2020) that may have affected how mothers interacted with their infants (Nicol-Harper et al., 2007) and how infants process faces (Bowman et al., 2021). Although the causal mechanism remains unclear, this is a unique and likely one-time dataset that could contribute to our understanding of how face identity processing changes in early development.

Our study has a number of other limitations. First, the sample size per group was small and spanned a wide age range. Our initial intention was to use this age variability to study perceptual narrowing. We combined data across ages to increase statistical power, knowing that face processing changes over this time (Pascalis et al., 2020). Additionally, although the groups were roughly matched (fortuitously) on key variables such as infant age, number of usable blocks, head motion, and eye gaze, the biological sex of the infants was not matched. There was an equal number of male and female infants post-lockdown, but all prelockdown infants were born female. The small sample size precludes any examination of sex differences. Although there are some advantages in face processing for female infants (Gluckman and Johnson, 2013), the evidence is mixed (Simpson et al., 2020; Maylott et al., 2021). Finally, although most data collection procedures were identical across groups, there was one key difference: whether the experimenters and parents wore face masks. It is possible that exposure to normal vs. obscured faces immediately prior to the experiment affected how infants processed faces. The serendipitous and opportunistic nature of this project means that we are saddled with these limitations, yet we believe the data remain valuable to report and may inform debates on the role of experience in the early development of face processing.

## Acknowledgements

We are thankful to the families of infants who participated. We also acknowledge the hard work of the Yale Baby School team, including L. Rait, J. Daniels, and K. Armstrong for recruitment, scheduling, and administration. Thank you to A. Klein, A. Bracher, and R. Blohowiak for help with gaze coding and to R. Watts for technical support. Thank you to the Beaver Brook Farm in Lyme, Connecticut for allowing us to photograph their sheep and to L. Rait and J. Wu for helping with stimulus creation. We are grateful for internal funding from the Department of Psychology and Faculty of Arts and Sciences at Yale University. T.S.Y was supported by an NSF Graduate Research Fellowship, and N.B.T-B. was further supported by the Canadian Institute for Advanced Research and the James S. McDonnell Foundation (https://doi.org/10.37717/2020-1208).

## Appendix

**Table A1.** Demographic information. The first six infants are the pre-lockdown group and the next six infants are the post-lockdown group. ‘ID’ is a unique infant identifier (i.e., sXXXX_Y_Z), with the first four digits (XXXX) indicating the family, the fifth digit (Y) the child number within family, and the sixth digit (Z) the session number with that child. ‘Age’ is recorded in months. ‘Sex’ is female or male. ‘TR prop’ is the proportion of TRs included from all usable blocks. ‘Total blocks’ is the number of blocks across all five conditions that were usable after motion and gaze exclusion. ‘Eye IRR’ is the proportion of frames coded the same way across gaze coders. ‘Days quarantine’ is the number of days between the start of the first lockdown in our State (March 15, 2020) and the day of the scan.

| ID | Age | Sex | TR prop | Total blocks | Eye IRR | Days quarantine |
| --- | --- | --- | --- | --- | --- | --- |
| s8687_1_5 | 17.1 | F | 1.000 | 15 | 0.976 | n/a |
| s6687_1_5 | 18.1 | F | 1.000 | 15 | 0.974 | n/a |
| s4607_1_4 | 13.0 | F | 0.982 | 10 | 0.929 | n/a |
| s4607_1_6 | 16.9 | F | 1.000 | 10 | 0.971 | n/a |
| s8037_1_2 | 12.2 | F | 0.976 | 15 | 0.934 | n/a |
| s7067_1_3 | 9.2 | F | 1.000 | 15 | 0.949 | n/a |
| s8687_2_1 | 10.7 | F | 0.975 | 14 | 0.935 | 244 |
| s6057_1_6 | 23.8 | M | 1.000 | 10 | 0.961 | 213 |
| s7047_1_2 | 17.6 | F | 1.000 | 15 | 0.958 | 216 |
| s3097_1_4 | 13.4 | F | 0.866 | 14 | 0.838 | 185 |
| s5477_1_1 | 13.3 | M | 0.933 | 15 | 0.956 | 215 |
| s5477_1_2 | 17.7 | M | 0.947 | 10 | 0.948 | 349 |

